# Cognitive training effects are shaped more by individual brain dynamics than age – Evidence from younger and older women

**DOI:** 10.1101/2025.07.23.666165

**Authors:** Zsófia Anna Gaál, Petia Kojouharova, Boglárka Nagy, Gwen van der Wijk, István Czigler, Andrea B. Protzner

## Abstract

Given the well-established structural and functional changes in the aging brain, it is widely assumed that cognitive aging is primarily driven by robust group-level differences between young and older adults. However, our individual-level EEG functional connectivity analysis challenges this notion. We investigated the impact of cognitive training on functional brain connectivity using task-switching paradigms in 39 younger (18–25 years) and 40 older (60–75 years) women. Participants were randomly assigned to either a training group, which completed eight individualized one-hour cognitive training sessions, or a no-contact control group. EEG was recorded at both pre- and post-training sessions across three task-switching paradigms (trained and near-transfer versions). Unique functional connectivity of different sources of variation was examined by calculating how much variance was shared across stable traits (e.g., individual, age, and common factors), or dynamic states (e.g., task and training effects).

Our results revealed that age accounted for only a modest proportion of variance, whereas self-similarity was a dominant factor – particularly in older adults. Similarly, group-level training effects were small but strongly modulated by individual neural profiles, suggesting person-specific trajectories. Participants recruited distinct neural networks across tasks, and even within the same task engaged unique, individual-specific network configurations, reflecting personalised brain adaptations to cognitive demands. Importantly, older adults displayed a shift from common to individual network patterns, consistent with increased neural specialization and compensatory mechanisms. These findings underscore the importance of moving beyond group-level contrasts toward models that capture the complexity of individual brain dynamics in cognitive aging and training responsiveness.

**Significance Statement:** Why do cognitive training programs work for some individuals but not for others? And is aging really the main factor driving changes in brain function? Using individual-level EEG functional connectivity analyses, we show that individual differences – not chronological age, task, or training – are the primary drivers of similarity in brain connectomes. While training effects are modest on average, their impact strongly depends on the individual’s unique neural profile. Older adults, in particular, rely more on personalized brain network configurations. These results help explain why cognitive training studies often yield inconsistent findings and point toward a more individualized approach to understanding and enhancing brain function across the lifespan.

## Introduction

Cognitive training refers to structured practice on tasks designed to enhance specific cognitive functions such as attention, memory, or executive control. These interventions aim to strengthen or maintain cognitive performance through repeated engagement with cognitively demanding activities. Identifying effective strategies to slow or prevent brain changes associated with brain aging, or to compensate for such changes is a central challenge in the field of cognitive aging and dementia research.

Despite extensive efforts, cognitive training has yielded mixed results. A major limitation has been the insufficient understanding of the mechanisms through which such interventions exert their effects (1,2). While training typically leads to improved performance on the practiced tasks, broader benefits have proven more elusive (3). Near-transfer effects – improvements on untrained tasks that rely on overlapping cognitive processes – are observed more consistently than far-transfer effects, which would reflect generalized gains in domains not directly targeted by training. However, even documented far-transfer effects tend to be modest in magnitude and of limited clinical relevance. Meta-analyses have reported inconsistent findings: some have shown benefits of cognitive training (4), while others have found no effect (5), fueling the debate over whether such gains transfer beyond the trained tasks (6, 2). Moreover, age may moderate training outcomes. While many studies report larger gains in younger than in older adults (e.g., 7, 8), which can be interpreted as a magnification effect due to greater cognitive resources, others have found larger benefits in children or older adults, consistent with compensation effects among individuals with lower baseline performance (e.g., 9, 2).

These heterogenous findings are further compounded by methodological variability across studies, including small sample sizes, differences in control conditions and session numbers, and the use of qualitatively different training protocols – such as strategy-based vs. process-based interventions (11). Previous work suggests that adaptive and multicomponent paradigms are generally more effective, although their implementation also varies across studies. Moreover, participants’ baseline cognitive abilities may also influence training outcomes: individuals with lower initial performance may benefit through compensatory mechanisms to bolster task-relevant skills, while higher-performing individuals may show a magnification of their existing abilities. For some tasks, there may be a threshold of performance below which participants are unlikely to benefit from training (11). Consequently, while some individuals exhibit substantial, generalizable gains that persist over time, others show little to no improvement.

We can approach this heterogeneity by focusing on the neural mechanisms that underlie training effects. One common view considers brain function as dynamic in that it reflects ongoing cognitive processes and can reconfigure with training. Indeed, previous work suggests that for brain structure, training-related changes include increases in grey matter and cortical volume (for a review, see 4). Functional changes are less consistent, but several studies have demonstrated decreased frontal and increased subcortical activation, suggesting that training may reduce the need for compensatory processes, especially for executive function. Precision neuroimaging (i.e., sampling sufficient data over multiple sessions to reveal meaningful individual divergence in the organisation of the brain) also has shown intra-individual change, such that functional networks can vary across tasks and time (e.g., 12, 2, 14). Alternatively, brain function can be viewed as stable in the sense that network properties reflect more stable traits such as age rather than cognitive content. For example, precision neuroimaging has shown that brain networks are relatively stable and show little variation over time and across states (e.g., 15, 2, 17, 18). In this latter case, brain function can be linked to long-term histories of regional co-activations. The truth likely lies along a continuum between these extremes, and it is therefore important to examine to what extent brain function reflects stable traits or dynamic states.

To do this, the present study leverages a previously published dataset that has already been analysed from both behavioural and electrophysiological perspectives, where the aim was to examine the extent, mechanisms, and durability of training-induced improvements and transfer effects in younger and older adults. In Gaál and Czigler (19), we examined performance and event-related potentials in reference and transfer task-switching paradigms. We found that after training, older adults reached the performance level of younger participants, with P3b components emerging after both cues and targets. Behavioural gains were also evident in near-transfer tasks and to a lesser extent in far-transfer ones, particularly in alerting and orienting networks. In Nagy et al. (20), we investigated changes in resting-state EEG following training, using spectral power density and multiscale entropy. The results revealed that cognitive training modulated age-related differences in resting-state dynamics, suggesting a shift toward more controlled and internally oriented processing in the older training group.

We now use this same dataset to examine stable and dynamic aspects of oscillatory activity during task performance. Previous work has characterized brain oscillations using power spectral density and functional connectivity (measured most commonly using phase-based metrics and envelope correlations; 21, 22) over different frequency bands. This is relevant given that distinct frequency bands have been associated with different cognitive functions. For example, in task-switching paradigms, delta oscillations are involved in inhibition and control processes (23, 24, 25). Theta oscillations are associated with flexible cognitive switching between task rules, memory retrieval and attentional control (26, 27, 28). Alpha oscillations reflect proactive control in task switching, through memory processes such as retrieving information from working- and long-term memory, and inhibitory processes involved in attention and response selection (23). Beta oscillations are involved in selective inhibition, decision-making, and memory processes (29, 30, 31). Age effects are most consistently observed in the theta and alpha bands. Theta activity, often associated with episodic and working memory processes, tends to increase at rest in older adults (32) or remain unchanged (33, 34), but may decline during task engagement (34, 35), possibly reflecting compensatory effort or reduced neural efficiency. Similarly, alpha power – particularly in the upper alpha range, is linked to attentional control and memory manipulation – typically decreases with age (32), which may reflect slower processing speed and reduced interference resolution (36). To better isolate age- and training-related neural effects, we focused on women – a group still underrepresented in the cognitive aging and training literature. This design choice also minimized variance associated with sex differences in brain structure, aging trajectories, and EEG signal characteristics. Building on this, we take a novel analytic approach to investigate the neural mechanisms underlying the observed training effects. Specifically, we examine the extent to which brain function reflects stable traits like individual, age, and common factors, or dynamic states such as task and training effects. By examining the relative contribution of these components, our aim is to move beyond group averages and toward a more nuanced understanding of person-specific brain dynamics in cognitive training.

## Materials and Methods

### Participants

Younger (ages 18–25) and older (ages 60–75) women were assigned to either a training (younger: n = 19, M = 21.4, SD = 1.7; older: n = 20, M = 65.3, SD = 3.3) or a control group (younger: n = 20, M = 21.7, SD = 1.6; older: n = 20, M = 66.1, SD = 3.1). All participants were right-handed, reported normal or corrected-to-normal vision, and had no history of neurological or psychiatric disorders. WAIS-IV (Wechsler Adult Intelligence Scale, 39) screening confirmed the absence of cognitive impairment in older adults (older control: M = 120.1, SD = 15.8; older training: M = 117.9, SD = 16.7). Participants provided written informed consent and were compensated for participation. The study was approved by the Joint Psychological Research Ethics Committee (Hungary).

### Procedure

Participants completed two sessions: Pre-training (baseline), and Post-training (after ∼1 month). WAIS-IV was administered at baseline and during subsequent sessions. EEG recordings were obtained in each session. The training group received eight 1-hour adaptive training sessions between Pre-training and Post-training; controls received no intervention.

Each EEG session included five tasks in fixed order: (1) resting-state (eyes open and closed); (2) informatively cued task-switching with nogo trials (reference task); (3) non-informatively cued task-switching with nogo trials (near-transfer); (4) informatively cued colour-shape classification (near-transfer); and (5) Attention Network Test (far-transfer) – tasks 2, 3 and 4 were analysed here. Instructions were delivered both orally and in writing. Stimuli were pseudorandomized and task order was consistent across sessions.

While deep phenotyping approaches benefit from the inclusion of multiple task paradigms, we restricted our analyses to three structurally similar task-switching tasks – the trained reference task and two near-transfer tasks (see Figure 1) – in which largest training-related effects have previously been demonstrated. This targeted selection enhances methodological consistency and allows us to better isolate the effects of cognitive training.

**Figure 1.**
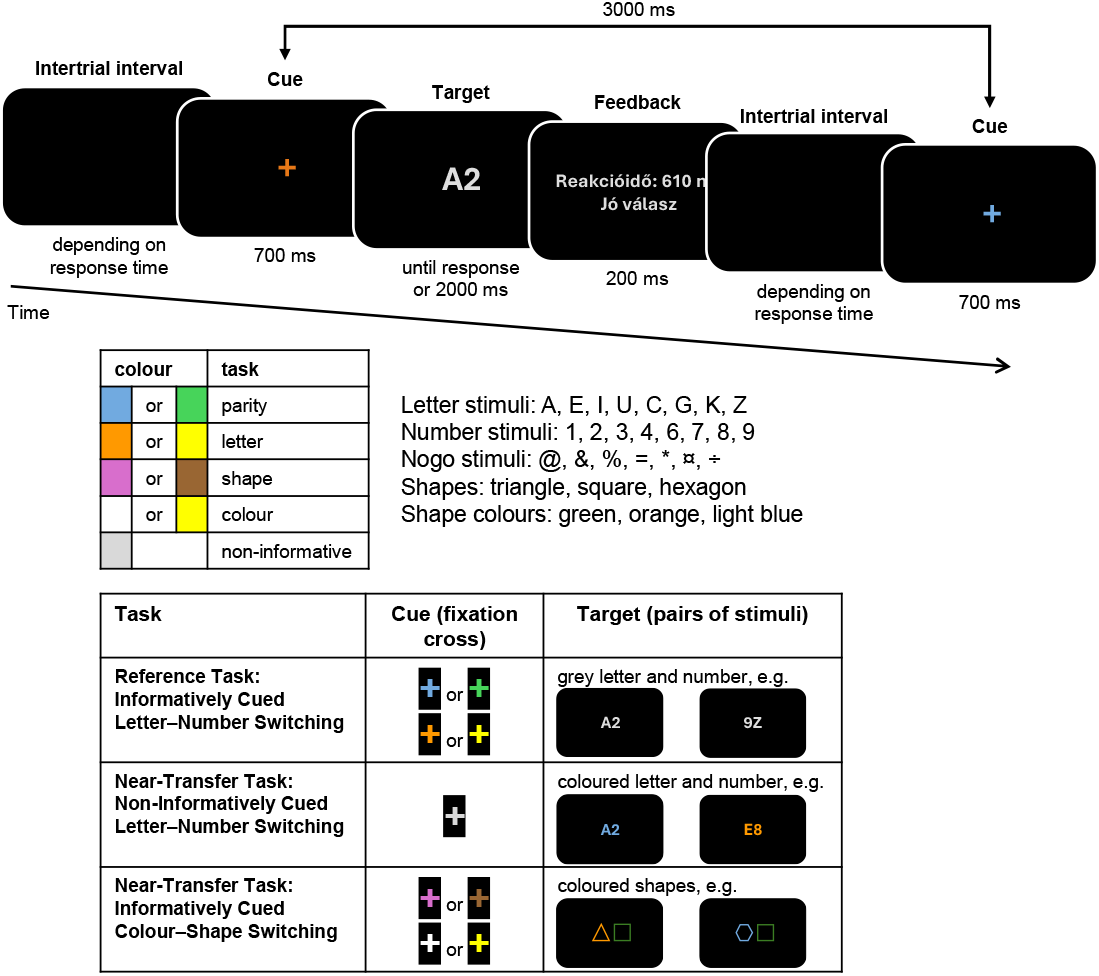
Schematic overview of the three task-switching paradigms. 1) Informatively cued task switching with nogo stimuli – letter classification and parity task; 2) non-informatively cued task switching with nogo stimuli – letter classification and parity task; and 3) informatively cued task switching – colour and shape classification task. Each trial began with a black screen, followed by a cue (informative in tasks 1 and 3, non-informative in task 2) displayed for 700 ms. The target then appeared (a letter-number pair with possible nogo stimuli in tasks 1 and 2; or a pair of shapes in task 3) and remained onscreen for 2,000 ms or until response. In the informatively cued conditions, cue colour indicated the relevant task. In the non-informatively cued condition, task identity was conveyed by the target’s colour. Feedback was provided for 200 ms following response.

### Reference Task: Informatively Cued Letter–Number Switching

This paradigm required participants to classify either a letter (vowel/consonant) or a number (odd/even) based on a preceding colour cue (700ms, yellow/orange for letter task; blue/green for the number task). Cues were never repeated consecutively to dissociate task switching from cue switching. Targets were grey letter-number pairs (1.8° x 1.4°), presented centrally for 2,000 ms or until response. Response mapping was counterbalanced across participants. In 25% of trials, a special character was presented, requiring participants to withhold their response (nogo trials). Both incongruently and congruently mapped targets were included to discourage strategic simplification; congruent trials and their immediate successors were excluded from analysis. Immediate feedback was provided after each trial for 200 ms. The task included single-task blocks (two each for letter and number classification; 104 trials per block) and mixed-task blocks (10 blocks; 500 trials total).

### Near-Transfer Task: Non-Informatively Cued Letter–Number Switching

Identical to the reference task except that cues were non-informative (grey), and task identity was conveyed solely by the colour of the target (yellow/orange for letter, blue/green for number). All other parameters matched the reference task.

### Near-Transfer Task: Informatively Cued Colour–Shape Switching

Participants classified pairs of geometric shapes (triangles, squares, hexagons; 5.5° x 2.75°) by either colour or shape based on the cue colour (purple/brown for shape; white/yellow for colour). Tasks included single-task blocks (1 per task, 48 trials/block) and mixed blocks (5 blocks, 240 trials total). No nogo trials were used in this paradigm.

### Training Session

Participants completed an informatively cued task-switching paradigm with nogo trials across 20 blocks of 50 trials each. Four tasks were cued by distinct colour pairs: (1) parity task (odd vs. even number), (2) letter classification task (vowel vs. consonant), (3) magnitude task (less than vs. greater than five), (4) case judgment task (uppercase vs. lowercase letter). Difficulty level was adjusted after each block based on performance: it increased following good performance, decreased after poor performance, and remained unchanged for intermediate outcomes.

#### EEG Recording and preprocessing

Participants were seated in an electrically and acoustically shielded room. Stimuli were presented using Presentation software at the centre of a monitor placed 125 cm from the participant. EEG was recorded at 1,000 Hz using NuAmps amplifiers (bandpass: DC–70 Hz) and NeuroScan 4.4 software (Compumedics, Victoria, Australia). Thirty-five passive Ag/AgCl electrodes were placed according to the 10–20 system at Fp1, Fp2, F7, F3, Fz, F4, F8, FT9, FC5, FC1, FC2, FC6, FT10, T7, C3, Cz, C4, T8, TP9, CP5, CP1, CP2, CP6, TP10, P7, P3, Pz, P4, P8, PO9, O1, Oz, O2, PO10. The reference was the tip of the nose, and FCz served as ground. Vertical and horizontal electrooculogram (EOG) signals were also recorded. All electrode impedances were kept below 10 kΩ.

Offline EEG data were processed in the following sequence: (1) the data were bandpass filtered using a finite impulse response (FIR) filter between 0.1 and 30 Hz (48dB/octave), (2) segmented into epochs time-locked to the target (from –100 ms to 1000 ms), (3) linear trends were removed, (4) baseline correction was applied using the prestimulus interval, and (5) trials exceeding ±80 μV on any channel were automatically rejected as artifacts. Additionally, the first trial of each block, error trials, congruently mapped target trials, and the trials immediately following them were excluded from further analysis.

#### Connectivity analysis

Using the Fieldtrip toolbox (40) in Matlab version 2019b, we estimated functional connectivity by calculating imaginary coherence (IMCOH; Nolte et al., 2004) across epochs for each unique pairing of 34 electrodes (561 pairs). IMCOH is a phase-based measure of the synchronization of brain regions at the EEG sensor level, and involves selecting only the imaginary component from a coherency calculation. It is robust to artifacts of volume conduction by being insensitive to zero time-lag synchrony between regions, thereby removing spurious connectivity.

#### Data analysis

We performed analyses on all groups together and each group separately to examine the similarity of functional connectomes matched on properties including participant, age, training, task and session. First, we extracted phase information for each epoch, channel, and frequency bin (ranging from 1 to 30 Hz, bins of 1 Hz) with Fieldtrip’s fast Fourier transformation algorithm. A Hanning taper was used. Then, we estimated functional connectivity for all electrode pairs within each individual, session, and task using IMCOH with Fieldtrip’s connectivity function. Next, the data were separated according to four frequency bands (delta: 1-4 Hz, theta: 4-7 Hz, alpha: 8-12 Hz, beta: 15-30 Hz), the IMCOH index was averaged for each frequency band, and the data were concatenated into a connectivity matrix for each frequency band, with averaged IMCOH values for each electrode pair, session, task, and individual (Supplementary Information Fig. S1b). In these matrices, each cell contained a connectivity (IMCOH) value, each column an electrode pair, and each row a level of each condition, e.g. the first row was the first individual’s first task and first session, the second row was the first individual’s first task and second session, and so on. Next, for each connectivity matrix we correlated the upper half of each session and individual’s whole-brain connectivity matrix across tasks with that of each other session (including sessions from the same individual) and each individual’s whole-brain connectivity pattern to construct a similarity matrix (Supplementary Information Fig. S1c). In this similarity matrix, each row and column represented a specific individual, task, and session. Fisher’s z-transformation was applied to these similarity matrices.

To quantify the magnitude of different sources of similarity, we calculated the average over different parts of the similarity matrix (as illustrated in Supplementary Information Fig. S2). These included all main effects: *Common* (all cells), *Session* (cells belonging to the same session), *Task* (cells belonging to the same task), *Age* (cells belonging to the same age group), and *Training* (cells belonging to the same training group). We also calculated average similarity for the intersections which represent all possible interactions between the conditions, e.g. *Session × Task, Age × Task*, and so on. To estimate the within-individual similarity, the cells belonging to the same individual were also averaged (*Individual*). Additionally, we calculated averages for the intersections *Individual × Session* and *Individual × Task* to analyse whether any differences in *Session* or *Task* are modified by individual characteristics. The main diagonal of the matrix (all ones by definition) was not included in the averages. In the separate-group analyses *Training, Age*, and their interactions were not included. The values were calculated for each participant and then submitted to dependent sample t-tests in MATLAB with permutation testing (1,000 permutations) to estimate significance. Specifically, for each comparison, the data were randomly shuffled between the two effects within an individual for each permutation. While the data were not fully independent, they were exchangeable, which is what is required for permutation testing (41). We compared each effect with its baseline (Supplementary Information Fig. S3b), i.e., each main effect to the *Common* effect, each interaction to the largest main effect included in it, and each higher-order interaction to the largest lower-order interaction. The *Individual* effect was compared to the largest main effect or interaction, while the *Individual × Session* and *Individual × Task* interactions were compared to the Individual effect. In other words, we tested whether the similarity in connectivity for each condition was significantly larger than its baseline. We used the false discovery rate (FDR) method to correct for multiple comparisons (42). We calculated the normalized relative effect magnitude as an indication of effect size. First, we subtracted the baseline from each effect (Supplementary Information Fig. S3a) and then divided by the sum of all baseline-corrected magnitudes. As such, normalized relative effect magnitudes reflect the proportion of each effect that was unique, i.e., not explained by its baseline, relative to the combined unique magnitude of all effects. If an effect had a lower magnitude than its baseline, the baseline-corrected magnitude was set to zero.

## Results

### Sources of similarity

We examined the contributions of different sources of similarity relative to their baselines across four frequency bands, both across all four groups combined and within each group separately.

### Combined analysis across for all four groups and three tasks

In this analysis, we focused solely on the effects of *Age*; other effects are addressed in separate group analyses. *Age* accounted for a significant portion of similarity across all frequency bands (1.7-5.2%). Additionally, the *Training × Age* (0.8-1.5%) and *Task × Age* (0.5-2.1%) interactions contributed significantly in all bands. *Session × Age* (0.5%) and the three-way *Training × Age × Session* (0.6%) interactions were significant only in the delta band, while the *Training × Age × Task* interaction was significant in both the delta and theta bands (1.4 and 0.7%). Nevertheless, the *Age* effects contributed a substantially smaller proportion to the similarity compared to the *Individual* FC (29.2–55.8%) and its interactions (1.0–19.3%), or the *Common* FC (22.6–38.5%).

**Figure 2.**
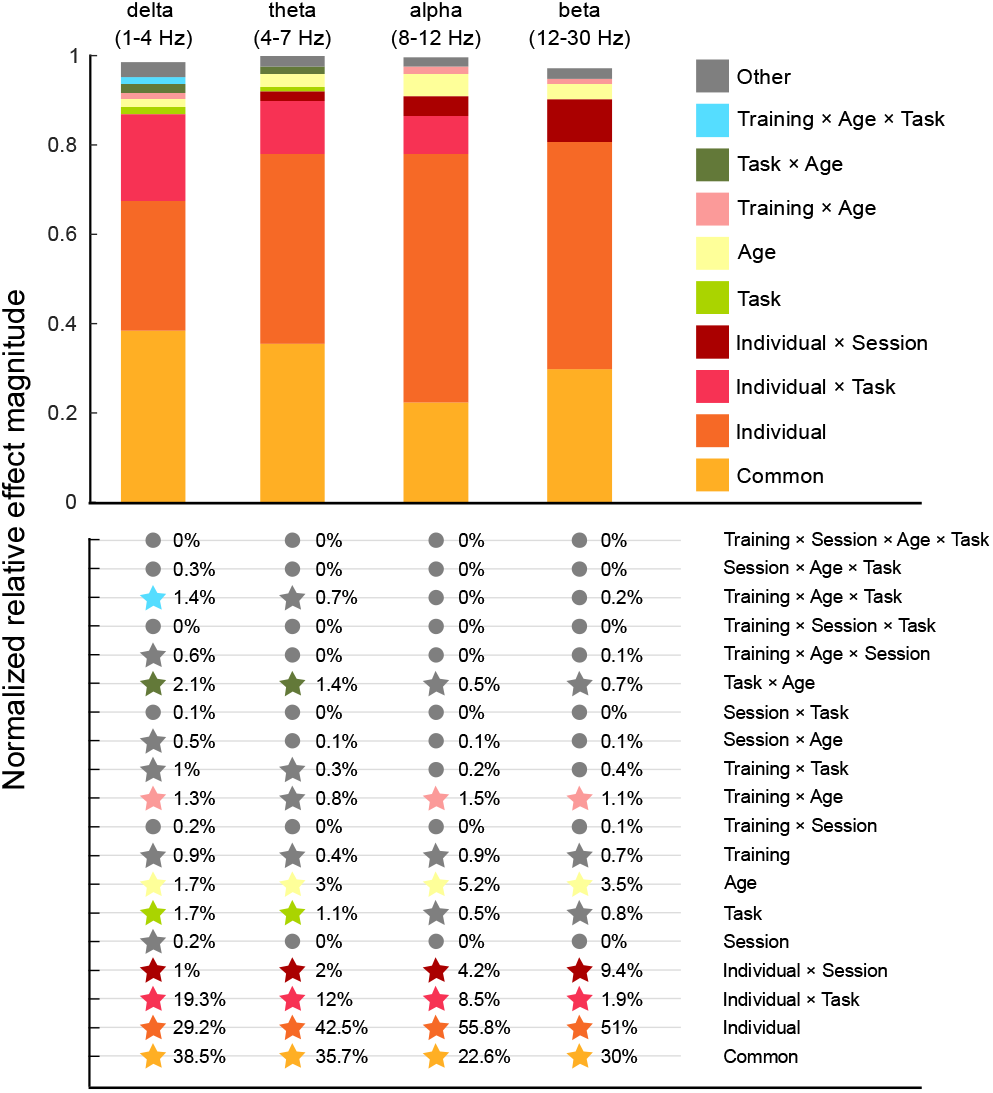
Normalized relative effect magnitude for each effect across all four groups and all frequency bands. Only significant effects are shown. Those explaining more than 1% of the similarity are displayed individually, while significant effects contributing less than 1% are grouped under ‘Other’. The percentage of explained similarity for each effect is shown below the bars. Stars indicate statistical significance. Although significant across all frequency bands, the contribution of *Age* to similarity was small compared to *Individual* and *Common* FC.

### Group-specific analyses across the three tasks

Session-specific FC contributed significantly to similarity only in the younger training group (delta, theta, and alpha bands, 0.4-3.5%), while the effect of training was reflected only in the *Individual × Session* interaction (theta, alpha, and beta bands, 3.7-8.6%) in the older training group. This interaction was also significant in the younger training group (delta, alpha, and beta bands, 9.3-11.4%), and in both control groups in the beta band, likely indicating practice effects in the latter. Task-specific FC had a small (0.52–3.00%) but significant contribution in all groups in the delta and theta bands. Importantly, in the control groups, the *Individual × Task* interaction was substantial in the delta, theta, and alpha bands (12.44–29.63%), indicating that individuals engaged distinct functional networks when performing different tasks. This interaction was smaller in the training groups – significant only in the theta band for younger adults and in the delta and theta bands for older adults – presumably because the *Individual × Session* interaction accounted for a larger proportion of similarity. Overall, *Individual* FC explained the largest proportion of similarity across the four groups (24.8–60.5%), with particularly high contributions in the older training group (37.8– 59.2%).

**Figure 3.**
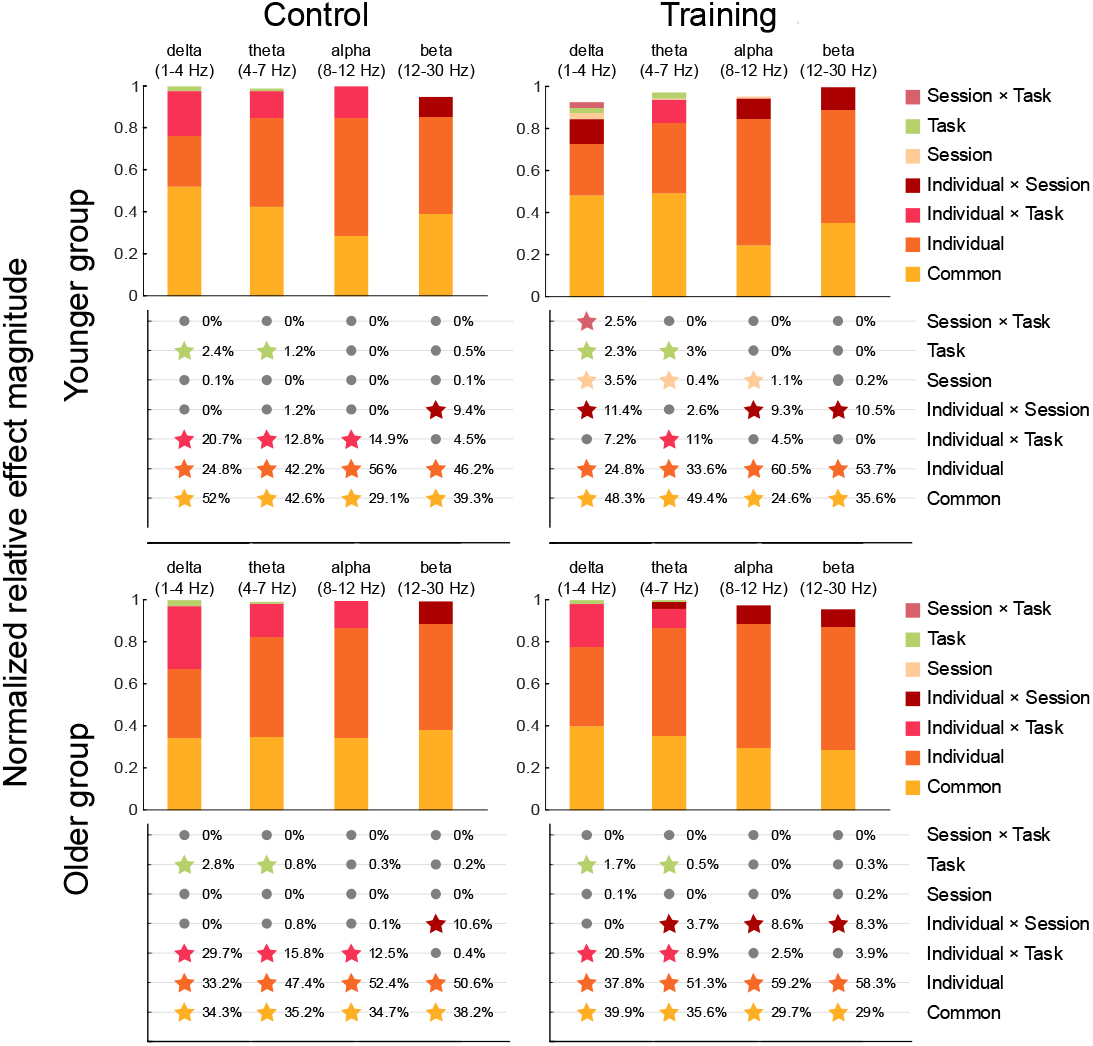
Normalized relative effect magnitude for each effect across frequency bands, computed separately for each of the four groups. Only significant effects are included in the bar graphs. The percentage of explained similarity by each effect is shown below the bars. Stars indicate statistical significance. The largest proportion of similarity was explained by Individual FC in the theta, alpha, and beta bands of older adults, and in the alpha and beta bands of younger adults. The second largest contribution came from the *Common* FC, which was most prominent in the delta band of older adults and in the delta and theta bands of younger adults. Together, these two factors accounted for 85.5–89.28% of similarity (normalized relative effect magnitude) in the beta band, and even in the delta band – where their contribution was the lowest – they still explained 67.54– 77.7%, indicating that task and training effects and their interactions were substantially smaller in comparison.

## Discussion

Building on our earlier work (19), we set out to identify the factors that shape cognitive training-induced changes in functional connectivity, with the broader goal of understanding how individual variability may inform the development of personalized interventions. Using a deep phenotyping approach (16, 17), we characterized the unique contributions of age, cognitive training and individual-level factors through EEG-based functional connectivity measures during task-switching paradigms in younger and older adults, who completed an eight-session adaptive training protocol designed to improve cognitive functions.

One of our most unexpected and intriguing findings was that age per se accounted for only a small proportion of the variance (1.75-5.18%) in EEG functional connectivity, despite the widespread assumption in the cognitive aging literature that young and older adults differ substantially. This suggests that individual differences may be more informative than chronological age when explaining variability in neural responses. Aging may reflect a shift toward increased interindividual variability rather than uniform decline.

In line with this age-associated shift, the main effect of *Session* (i.e., pre–post change) was not significant in the older training group – similarly to what was expected and observed in both control groups – but the *Individual × Session* interaction accounted for substantial similarity. This indicates that training responses in older adults are highly individualized, and that group-level effects may underestimate the functional plasticity retained in later life. In contrast, younger adults showed a significant main effect of *Session* in all frequency bands except beta, yet even here, the *Individual × Session* interaction explained a much larger share of the similarity (except in the theta band), again underscoring the dominance of idiosyncratic training trajectories. Importantly, in both age groups, and especially in the training conditions, the *Individual × Session* interaction was substantially larger than the main effect of *Session*, indicating that the effectiveness of cognitive training cannot be fully captured by group-average changes. Instead, training appears to interact strongly with individual neural architectures or strategies, potentially explaining the high variability across cognitive training studies in the literature. These findings are consistent with the revised Scaffolding Theory of Aging and Cognition (STAC-r, 37), suggesting that functional reorganization and compensatory mechanisms – such as functional redundancy – may underlie training responsiveness in older age.

In addition, we observed a robust *Individual × Task* interaction, more pronounced in the control group, indicating that participants – including those in the control condition – engaged distinct neural networks to solve different tasks, even within the same experimental condition. Thus, we found no evidence in favour of a single, unified neural response pattern, but rather identified multiple connectivity patterns that vary across individuals. Interestingly, training appeared to reduce this heterogeneity, possibly by promoting more efficient or convergent network recruitment. A significant *Task × Session* interaction further supports the notion that training altered functional connectomes, albeit secondarily to the more nuanced individual-level patterns.

Finally, we observed that the proportion of explained similarity attributable to *Common* vs. *Individual* components differed across age groups: in younger adults, *Common* variance dominated, whereas in older adults, *Individual* variance was more prominent. This pattern suggests that network specialization increases with age, possibly reflecting the ways that an individual’s unique experiences interact with brain aging processes.

Most of our significant effects were observed across all frequency bands, indicating that training and task-related changes broadly influence neural networks. Exceptions to this pattern were the *Task* and *Task × Individual* effects, which were most prominent in the delta and theta bands. This may reflect the task-specific demands of the paradigm including inhibitory and control processes, which have been linked to delta band activity (23, 24, 25), and cognitive switching and attentional control which has been linked to theta activity (26, 28).

By focusing on women, our study offers new insights into age and training-related connectivity effects. However, the relatively small and WEIRD (Western, Educated, Industrialized, Rich, and Democratic, 43) sample limits generalizability. Future studies should test whether the findings extend across genders, socioeconomic backgrounds, and clinical populations.

Taken together, our findings highlight that chronological age is not the primary determinant of training-related functional changes. Rather, they underscore the central role of individual-level variance in shaping how cognitive training effects unfold in the brain. We emphasize that, in most of the cognitive training literature, references to individual differences usually pertain to demographic characteristics, socio-economic status, motivation, personality traits or baseline cognitive performance (38). In our study, however, we focus on a different level of individuality – specifically, the uniqueness and stability of each participant’s functional brain connectome. This neural individuality appears to strongly influence both the magnitude and the trajectory of training related brain responses, suggesting that truly effective and transferable interventions will need to move beyond group averages and adopt personalized approaches. In this context, understanding the functional architecture of individual brains may offer a promising route toward more targeted, efficient, and impactful cognitive training programs across the lifespan.

## Acknowledgments

We thank Emese Várkonyi for her technical assistance. The research was supported by the Hungarian Research Fund (OTKA PD 101175 and OTKA K 145940) and the János Bolyai Research Scholarship of the Hungarian Academy of Sciences to ZAG, and by a Natural Sciences and Engineering Research Council of Canada Discovery Grant (05299–2020) to ABP.

