## Supplementary material for "Cognitive training effects are shaped more by individual brain dynamics than age – Evidence from younger and older women": SI Appendix

**This PDF file includes:**

Figures S1 to S3

SI References

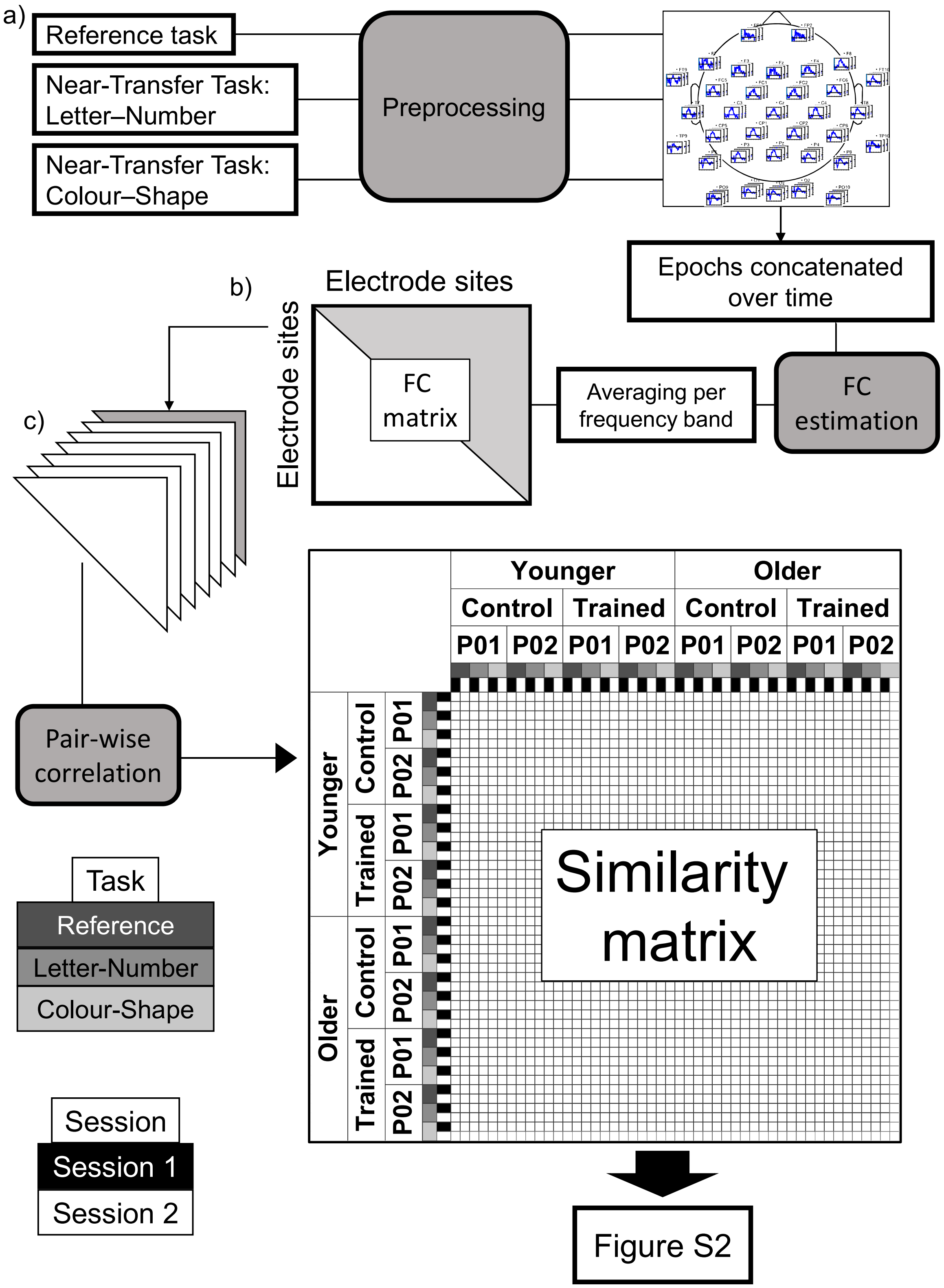


**Fig. S1.** Overview of the analysis procedure. The example shown illustrates the analysis including all groups, but the same steps were applied in the group-specific analyses. a) The EEG data for the target stimuli from the Informatively Cued Letter–Number Switching (reference), Non-Informatively Cued Letter–Number Switching (near-transfer), and Informatively Cued Colour–Shape Switching (near-transfer) tasks were preprocessed using the same pipeline and concatenated separately for each participant, task, and session. b) Functional connectivity (FC) was then calculated as the correlation between the time courses of each electrode pair for each frequency from 1 to 30 Hz, yielding an electrode-by-electrode FC matrix for each participant, task, session, and frequency. These matrices were then averaged for four frequency bands: delta (1-4 Hz), theta (4-7 Hz), alpha (8-12 Hz), and beta (12-30 Hz). c) The upper triangle (grey) of each FC matrix was correlated with all others (across participants, tasks, and sessions) to generate a similarity matrix. This procedure was conducted separately for each frequency band, resulting in four similarity matrices.

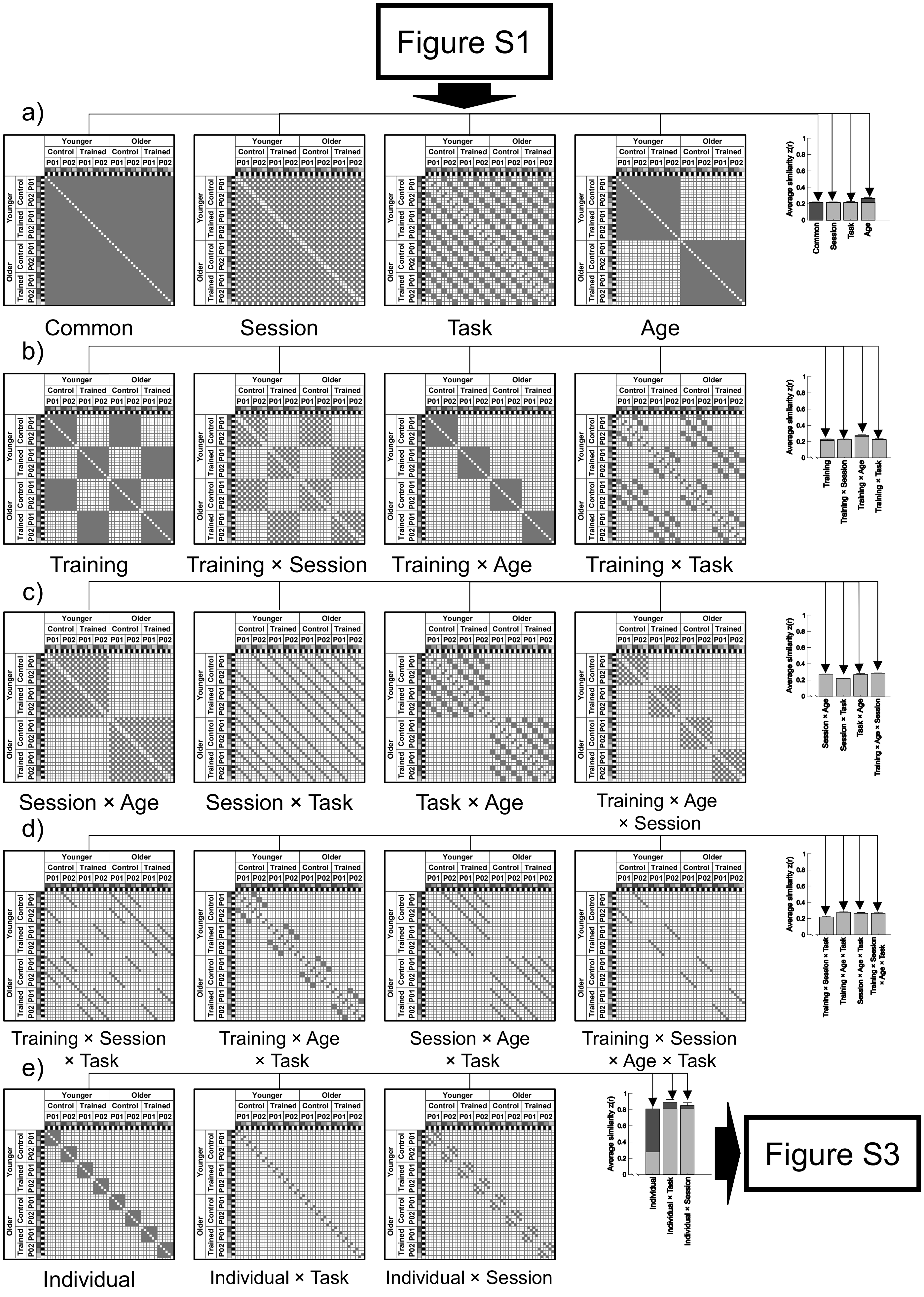
Figure S1
a)
Younger
Older
Control
Trained
P01
P02
Average similarity z(r)
Common
Session
Task
Age
b)
Training
Training × Session
Training × Age
Training × Task
c)
Session × Age
Session × Task
Task × Age
Training × Age × Session
d)
Training × Session × Task
Training × Age × Task
Session × Age × Task
Training × Session × Age × Task
e)
Individual
Individual × Task
Individual × Session
Figure S3

**Fig. S2.** Schematic overview of different sources of similarity in the data. The analysis including all groups is used as an example, but the same steps were applied in the group-specific analyses. The contribution of each source of variance was estimated by calculating average similarity over different cell configurations of the similarity matrix. The presented patterns display the configuration for the calculation in the following order: a) **Row 1** (left to right): *Common effect* (similarities in FC across all participants and sessions), *Session effect* (similar FC across participants within time points), *Task effect* (similarities in FC across participants within tasks), *Age effect* (similarities in FC among older and younger participants), b) **Row 2** (left to right): *Training effect* (similarities in FC among trained participants and controls), *Training × Session interaction* (similarities in FC among trained participants and controls within time points), the *Training × Age interaction* (similarities in FC among older trained participants, younger trained participants, older controls, and younger controls), *Training × Task interaction* (similarities in FC among trained participants and controls within tasks), c) **Row 3** (left to right): *Session × Age interaction* (similarities in FC among older and younger participants within time points), *Session × Task interaction* (similarities in FC across participants within time points and tasks), *Task × Age interaction* (similarities in FC among older and younger participants within tasks), *Training × Age × Session interaction* (similarities in FC among older trained participants, younger trained participants, older controls, and younger controls within time points); d) **Row 4** (left to right): *Training × Session × Task interaction* (similarities in FC among trained participants and controls within time points and tasks), *Training × Age × Task interaction* (similarities in FC among older trained participants , younger trained participants, older controls, and younger controls within tasks), *Session × Age × Task interaction* (similarities in FC among older and younger participants within time points and tasks), *Training × Session × Age × Task interaction* (similarities in FC among older trained participants, younger trained participants, older controls, and younger controls within time points and tasks), and e) **Row 5** (left to right): *Individual effect* (similarities in FC within individuals), *Individual × Task interaction* (similarities in FC within individuals and tasks), *Individual × Session interaction* (similarities in FC within individuals and time points).

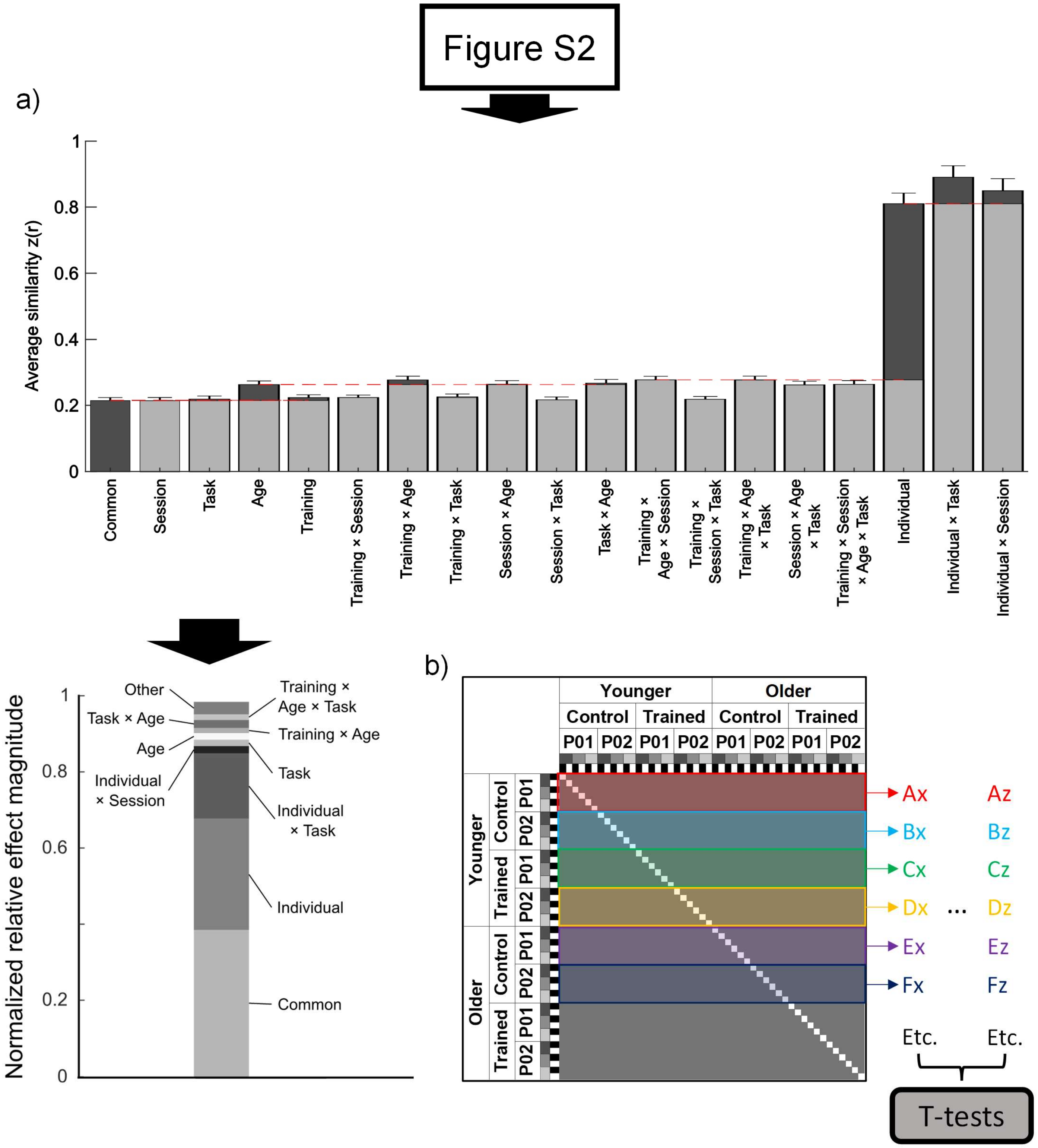


**Fig. S3.** Calculation of effect magnitudes. The analysis including all groups is used as an example, but the same steps were applied in the group-specific analyses. a) From the average similarity for each effect, we calculated the normalized relative effect magnitude by first subtracting the baseline for each effect (indicated by the dashed red lines) and then dividing by the total explained similarity. If an effect was smaller than its baseline, its magnitude was set to zero. b) We calculated the average similarity for each effect per individual by taking the average of the patterns for the six rows representing each participant (example shown for the common effect, where each row represents one participant's similarity to all other participants across tasks and session). We then used dependent samples t-tests to test whether these participant-specific similarities contributed significantly to the overall similarity structure of the data.

Figures S1, S2, and S3 adapted from Figure 1 in van der Wijk et al. (2024).
